# MAIT cells are minimally responsive to Mycobacterium tuberculosis within granulomas, but are functionally impaired by SIV in a macaque model of SIV and Mtb co-infection

**DOI:** 10.1101/2020.01.07.897447

**Authors:** Amy L. Ellis-Connell, Alexis J. Balgeman, Erica C. Larson, Mark A. Rodgers, Cassaundra Ameel, Tonilynn Baranowski, Pauline Maiello, Jennifer A. Juno, Charles A. Scanga, Shelby L. O’Connor

## Abstract

Mucosal associated invariant T (MAIT) cells recognize and can directly destroy bacterially infected cells. While a role for MAIT cells has been suggested in several *in vitro* and in vivo models of *M.tuberculosis* (Mtb) infection, these studies have often focused on MAIT cells within the peripheral blood or are cross-sectional studies rather than longitudinal studies. The role of MAIT cells within granulomas and other sites of Mtb infection is relatively unknown. Furthermore, how HIV/SIV infection might impair MAIT cells at the sites of Mtb infection has not been determined. Using a Mauritian cynomolgus macaque (MCM) model system, we phenotyped MAIT cells in the peripheral blood and BAL prior to and during infection with SIVmac239. To characterize the role of MAIT cells within granulomas, SIV+ and -naïve MCM were infected with a low dose of Mtb for 6 weeks. MAIT cell frequency and function was examined within the peripheral blood, distal airways, as well as within Mtb-affected lymph nodes (LN) and tissues. Surprisingly, we found no evidence of MAIT cell responsiveness to Mtb within granulomas. Additionally, MAIT cells only minimally responded to mycobacterial stimulus in ex vivo functional assays. In contrast, most MAIT cell activation seemed to occur in samples with highly active SIV replication, including blood and SIV-infected LN. Finally, the ability of MAIT cells to secrete TNFα (TNF) was impaired during SIV and Mtb co-infection, indicating that the two pathogens together could have a synergistically deleterious effect on MAIT cell function. The effect of this functional impairment on overall TB disease burden was unclear, but might be deleterious if MAIT cells are needed to fully activate antimycobacterial immune cells within the granulomas.

## Introduction

*Mycobacterium tuberculosis* (Mtb) is the causative agent of tuberculosis (TB), and 10 million new Mtb infections occurred in 2018 alone (https://www.who.int/tb/publications/global_report/en/). Ninety (90) percent of healthy humans are able to immunologically control Mtb infection; however, TB remains a major global health concern. One factor that can complicate the outcome of Mtb infection is Human Immunodeficiency Virus (HIV) co-infection. HIV+ individuals are 20 times more likely to develop active TB disease and Mtb infection is the number one cause of morbidity and mortality in HIV+ individuals (https://www.who.int/hiv/mediacentre/news/hiv-tb-patient-centred-care/en/).

The immune correlates of protection from Mtb that are lost in HIV+ individuals are poorly understood, particularly within the granulomas and affected tissues themselves [1]. The adaptive immune system, particularly CD4+ T lymphocytes, is important for containing Mtb within granulomas. While these cells play key roles in Mtb containment and are impaired in frequency and function during HIV infection, they are likely not the sole correlate of protection during Mtb infection.

One understudied population of cells with regards to its role in Mtb infection are mucosal associated invariant T (MAIT) cells. MAIT cells are innate immune T lymphocytic cells that recognize metabolites from bacterial antigens [2–5]. Previous studies, both *in vitro and in vivo*, have suggested a potential role for MAIT cells in Mtb infection. In humans as well as in macaques, MAIT cells are highly abundant in the lungs and bronchoalveolar lavage fluid, and express activation markers such as CD69 in the blood following Mtb infection [6,7]. In humans, MAIT cell frequencies are altered in the blood during active TB [8–10]. Several *in vitro* studies have demonstrated the ability of MAIT cells to respond to antimycobacterial antigens [3]. Furthermore, knocking out the major histocompatibility-related receptor 1 (MR1) receptor, which displays bacterial metabolites to MAIT cells, results in exacerbated TB in mice [5]. However, a new report performed in Mtb-infected rhesus macaques indicated that MAIT cells may not respond to Mtb infection within the tissues or within the airways [11]. Indeed, little evidence exists that MAIT cells respond to Mtb within the granulomas and affected LN.

Conflicting evidence also exists regarding the impairment of MAIT cells during HIV infection. Several cross-sectional human studies indicate that MAIT cells are depleted during HIV infection[12,13] [8,10,12,14]. In agreement with these studies, MAIT cell frequency and function were altered in a cross-sectional study during SIV infection of rhesus macaques [15]. One caveat for these human and macaque studies is that many of these measured MAIT cells in the blood only. Furthermore, these studies were not performed longitudinally within individuals. A recent report examining MAIT cells within the lungs of HIV+ and HIV-naïve patients with active TB indicated that MAIT cell frequency was actually higher in the lungs of the HIV+ patients compared to the HIV-naïve patients [16]. Additionally, a recent study indicated that in SIV-infected pigtailed macaques, MAIT cell frequency and function were not impaired longitudinally within the blood [17]. Overall, few studies to date have focused on how HIV/SIV infection affects MAIT cell frequency and/or function within granulomas of Mtb-infected individuals.

Our laboratory previously established a macaque model of Mtb and SIV co-infection to study how pre-existing SIV infection impairs the adaptive immune response to Mtb. We used Mauritian cynomolgus macaques (MCM), which have simplified major histocompatibility complex (MHC) genetics, for these studies. SIV-positive or -naïve MCM were infected with a low dose of Mtb for 12-20 weeks. While no difference in TB disease between the two cohorts was observed at 4 weeks post-Mtb infection [18], at 8 weeks the SIV co-infected macaques had significantly more granulomas than the SIV-naïve macaques [18]. Overall, the TB disease burden within the lungs of the co-infected macaques was greater than in the SIV-naïve macaques at study endpoint [18].

Building upon our previous study, we hypothesized that MAIT cells in Mtb-infected granulomas and LN during early Mtb infection might function as early responders to Mtb. Furthermore, we hypothesized that pre-existing SIV infection might impair the frequency and/or function of MAIT cells within granulomas and affected LN. To test these hypotheses, MCM were first infected SIVmac239 for 6 months followed by co-infection with a low dose of Mtb for 6 weeks. The control group was infected with Mtb only. There were no striking clinical differences between the two groups at necropsy just 6 weeks post Mtb. Differences in the adaptive immune response within granulomas and affected LN consisted of higher expression of activation/exhaustion markers and lower TNF production by T cells (Larson et. al., manuscript in prep). Here, MAIT cells were fully characterized longitudinally during a 6-month SIV infection to test whether SIV impaired their frequency or function. We also assessed MAIT cell frequency and function during Mtb infection in blood, BAL, and Mtb-affected tissues of SIV-naïve and SIV-infected MCM.

## MATERIALS AND METHODS

### Animal care

Mauritian cynomolgus macaques (*macaca fascicularis*; MCM) were obtained from Bioculture, Ltd. (Mauritius). MCM with at least one copy of the M1 MHC haplotype were chosen for this study. The macaques (n=4) infected with SIV alone were cared for by the staff at the Wisconsin National Primate Research Center (WNPRC) in accordance with the regulations, guidelines, and recommendations outlined in the Animal Welfare Act, the Guide for the Care and Use of Laboratory Animals, and the Weatherall Report. The University of Wisconsin-Madison (UW-Madison), College of Letters and Science and Vice Chancellor for Research and Graduate Education Centers Institutional Animal Care and Use Committee approved the nonhuman primate research covered under protocol G005507. The University of Wisconsin-Madison Institutional Biosafety Committee approved this work under protocol B00000205. All macaques were housed in standard stainless steel primate enclosures providing required floor space and fed using a nutritional plan based on recommendations published by the National Research Council. Macaques had visual and auditory contact with each other in the same room. Housing rooms were maintained at 65– 75°F, 30–70% humidity, and on a 12:12 light-dark cycle (ON: 0600, OFF: 1800). Animals were fed twice daily a fixed formula, extruded dry diet with adequate carbohydrate, energy, fat, fiber, mineral, protein, and vitamin content (Harlan Teklad #2050, 20% protein Primate Diet, Madison, WI) supplemented with fruits, vegetables, and other edible objects (e.g., nuts, cereals, seed mixtures, yogurt, peanut butter, popcorn, marshmallows, etc.) to provide variety to the diet and to inspire species-specific behaviors such as foraging. To further promote psychological well-being, animals were provided with food enrichment, structural enrichment, and/or manipulanda. Environmental enrichment objects were selected to minimize chances of pathogen transmission from one animal to another and from animals to care staff. While on study, all animals were evaluated by trained animal care staff at least twice daily for signs of pain, distress, and illness by observing appetite, stool quality, activity level, physical condition. Animals exhibiting abnormal presentation for any of these clinical parameters were provided appropriate care by attending veterinarians. Prior to all minor/brief experimental procedures, macaques were anesthetized with an intramuscular dose of ketamine (10 mg kg^-1^) and monitored regularly until fully recovered from anesthesia. Per WNPRC standard operating procedure (SOP), all animals received environmental enhancement included constant visual, auditory, and olfactory contact with conspecifics, the provision of feeding devices which inspire foraging behavior, the provision and rotation of novel manipulanda (e.g., Kong toys, nylabones, etc.), and enclosure furniture (i.e., perches, shelves). At the end of the study, euthanasia was performed following WNPRC SOP as determined by the attending veterinarian and consistent with the recommendations of the Panel on Euthanasia of the American Veterinary Medical Association. Following sedation with ketamine (at least 15mg/kg body weight, IM), animals were administered at least 50 mg/kg IV or intracardiac sodium pentobarbital, or equivalent, as determined by a veterinarian. Death was defined by stoppage of the heart, as determined by a qualified and experienced individual.

MCM at the University of Pittsburgh (U.Pitt., n=19) were housed in a BSL2+ animal facility during SIV infection and then moved into a BSL3+ facility within the Regional Biocontainment Laboratory for infection with *Mtb*. Animal protocols and procedures were approved by the U.Pitt. Institutional Animal Care and Use Committee (IACUC) which adheres to guidelines established in the Animal Welfare Act and the Guide for the Care and Use of Laboratory Animals (8th Edition). The U.Pitt. IACUC reviewed and approved the study protocols 18032418 and 15035401, under Assurance Number A3187-01). The IACUC adheres to national guidelines established in the Animal Welfare Act (7 U.S.C. Sections 2131–2159) and the Guide for the Care and Use of Laboratory Animals (8th Edition) as mandated by the U.S. Public Health Service Policy. Macaques were housed at U.Pitt. in rooms with autonomously controlled temperature, humidity, and lighting. Animals were pair-housed in caging measuring 4.3 square feet per animal and spaced to allow visual and tactile contact with neighboring conspecifics. The macaques were fed twice daily with biscuits formulated for nonhuman primates, supplemented at least 4 days/week with fresh fruits, vegetables or other foraging mix. Animals had access to water *ad libitem*. An enhanced enrichment plan is designed and overseen by our nonhuman primate enrichment specialist. This plan has three components. First, species-specific behaviors are encouraged. All animals have access to toys and other manipulanda, some of which are filled with food treats (e.g. frozen fruit, peanut butter, etc.). These are rotated on a regular basis. Puzzle feeders, foraging boards, and cardboard tubes containing small food items also are placed in the cage to stimulate foraging behaviors. Adjustable mirrors accessible to the animals stimulate interaction between cages. Second, routine interaction between humans and macaques are encouraged. These interactions occur daily and consist mainly of small food objects offered as enrichment and adhere to established safety protocols. Animal caretakers are encouraged to interact with the animals by talking or with facial expressions while performing tasks in the housing area. Routine procedures (e.g. feeding, cage cleaning, etc.) are done on a strict schedule to allow the animals to acclimate to a routine daily schedule. Third, all macaques are provided with a variety of visual and auditory stimulation. Housing areas contain either radios or TV/video equipment that play cartoons or other formats designed for children for at least 3 hours each day. The videos and radios are rotated between animal rooms so that the same enrichment is not played repetitively for the same group of animals. All animals are checked at least twice daily to assess appetite, attitude, activity level, hydration status, etc. Following SIV and/or Mtb infection, the animals are monitored closely for evidence of disease (e.g., anorexia, lethargy, tachypnea, dyspnea, coughing). Physical exams, including weights, are performed on a regular basis. Animals are sedated prior to all veterinary procedures (e.g. blood draws, etc.) using ketamine or other approved drugs. Regular PET/CT imaging is conducted on our macaques following Mtb infection and has proved very useful for monitoring disease progression. Our veterinary technicians monitor animals especially closely for any signs of pain or distress. If any are noted, appropriate supportive care (e.g. dietary supplementation, rehydration) and medications (e.g. analgesics) are given. Any animal considered to have advanced disease or intractable pain or distress from any cause is sedated with ketamine and then humanely euthanized using sodium pentobarbital (65 mg/kg, IV). Death is confirmed by lack of both heartbeat and pupillary responses by a trained veterinary professional.

### SIV and Mtb infection of MCM

At UW-Madison, 4 MCMs were infected intrarectally with 3,000 TCID50 SIVmac239. Six months after infection, animals were humanely euthanized as indicated in the section above.

For studies performed at U.Pitt., animals in the SIV/Mtb co-infection group (n=8) were infected intrarectally with 3,000 TCID_50_ SIVmac239. After 6 months, the animals were co-infected with a low dose (3-12 CFU) of Mtb (Erdman strain) via bronchoscopic instillation, as described previously [18]. Animals in the SIV-naïve control group (n=11) were infected with Mtb in an identical manner. TB progression was monitored by clinical testing and PT/CT imaging (Larson et al., manuscript in prep). Six weeks after Mtb infection, animals were humanely euthanized and necropsies were performed using PET/CT imaging to guide the excision of all granulomas. Random samples from each lung lobe were also harvested, as well as peripheral (axillary and inguinal) LN, all thoracic LN, mesenteric LN, liver, and spleen ([19];further described in Larson et al., manuscript in prep).

### Sample collection

For all cohorts, whole blood and plasma samples were collected longitudinally pre- and post-SIV and/or Mtb infection, and peripheral blood mononuclear cells (PBMC) and were isolated by Ficoll density gradient purification as previously described [18,20]. Bronchoalveolar lavage (BAL) samples were also collected at the indicated time points pre- and post-SIV and/or Mtb infection as previously described [18].

Tissue samples were collected at necropsy, as detailed above. For the SIV-only cohort, peripheral and thoracic LN were collected, single cell suspensions were prepared, and samples were stained for flow cytometry. For the SIV-naïve and SIV+ MCM infected with Mtb at U.Pitt, the tissue samples listed above were homogenized using Medimachines (BD Biosciences) and each homogenates was divided. One portion was plated on 7H11 agar to quantify the Mtb colony forming units (CFU) as previously described ([19] [18], and the other portion was used for flow cytometry.

### Plasma viral loads and virus sequencing

Plasma viral loads were quantified as previously described [20–23]. Briefly, viral RNA was isolated from plasma, reverse transcribed, and amplified with the SuperScript III Platinum one-step quantitative RT-PCR system (Thermo Fisher Scientific). Samples were then run on a LightCycler480 (Roche) and compared to an internal standard curve on each run.

### Flow Cytometry

To characterize MAIT cells from peripheral blood, BAL, LN, and granulomas, the rhesus macaque (Mamu) MR1 tetramer loaded with either the 5-OP-RU active metabolite or the Ac-6-FP control metabolite were used [24]. The MR1 tetramer technology was developed jointly by Dr. James McCluskey, Dr. Jamie Rossjohn, and Dr. David Fairlie, and the material was produced by the NIH Tetramer Core Facility as permitted to be distributed by the University of Melbourne. For PBMC, staining was performed using cryopreserved cells collected longitudinally from the current study, or from cryopreserved cells from SIV-naïve MCM collected from previous, unrelated studies. For BAL and tissue samples, freshly isolated cell homogenates were used for staining.

Approximately 1×10^6^ cells (or as many cells as possible from granulomas with fewer than1×10^6^ cells) were stained with 0.25 ug of Mamu MR1 5-OP-RU or Ac-6-FP tetramer for one hour in the presence of 500 nM Dasatinib (Thermo Fisher Scientific; Cat No. NC0897653). When TCRVα7.2 co-staining was performed, the antibody was added 30 minutes after the addition of the MR1 tetramer. Cells were washed once with FACS buffer (10% Fetal Bovine Serum (FBS) in a 1X PBS solution) supplemented with 500 nM Dasatinib, then surface antibody staining was performed for 20 minutes in FACS buffer + 500 nM Dasatinib. A complete list of antibodies used for surface staining is shown in Table 1. Samples were fixed in 1% paraformaldehyde for a minimum of 20 minutes. For intracellular staining, cells were washed twice with FACS buffer and staining with antibodies was performed in Medium B permeabilization buffer (Thermo Fisher Scientific, Cat. No. GAS002S-100) for 20 minutes at room temperature. For intranuclear staining, the True Nuclear Transcription Factor Buffer Set (Biolegend; San Diego, CA) was used according to manufacturer’s instructions. Briefly, cells were fixed in the TrueNuclear™ fixation solution for 1 hour, then washed three times with the permeabilization buffer. Cells were then stained with the transcription factors indicated in Table 1 at 4°C for one hour, rinsed three times with permeabilization buffer, then resuspended in FACS buffer. Flow cytometry was performed on a BD LSR II (Becton Dickinson; Franklin Lakes, NJ), and the data were analyzed using FlowJo software for Macintosh (version 9.9.3 or version 10.1).

**Table 1.** Antibodies used in staining panels for flow cytometry.

| Marker | Clone(s) | Fluorochrome(s) | Purpose of Use | Surface/Intracellular/intranuclear |
| --- | --- | --- | --- | --- |
| CD45 | D058-1283 | BV786 | Lineage | Surface |
| CD3 | SP34-2 | AF700, BV650 | T cell marker | Surface |
| CD4 | OKT4, L200 | PE Cy7, BV510, BV711, BV786 | T cell marker | Surface |
| CD8 | RPAT8, SK1 | PacBlue, BV510, BV605, BV711, BV786 | T cell marker | Surface |
| CD206 | 19.2 | PE Cy5 | Macrophage exclusion | Surface |
| TCRVα7.2 | 3C10 | BV421, BV605 | MAIT cell marker | Surface |
| CCR5 | J418F1 | BV421 | Chemokine/MAIT lung homing | Surface |
| CCR6 | 11A9 | PE CF594, PE Cy7 | Chemokine/gut homing | Surface |
| CD127 | MB15-18C9 | PE | IL-7 receptor | Surface |
| CXCR3 | G025H7 | PE Dazzle 594 | Chemokine/MAIT lung homing | Surface |
| CCR7 | FAB197F | FITC | Chemokine/lymph homing | Surface |
| CD28 | CD28.2 | APC, BV510 | T cell memory | Surface |
| CD95 | DX2 | PE Cy5 | T cell memory | Surface |
| T-Bet | 4B10 | BV605 | Transcriptional marker | Intranuclear |
| Eomes | WD1928 | PE Cy7 | Transcriptional marker | Intranuclear |
| RORγT | AFKJS-9 | PE | Transcriptional marker | Intranuclear |
| PLZF1 | R17-809 | PE CF594 | Transcriptional marker | Intranuclear |
| Fox3P | 150D | AF488, AF647 | Transcriptional marker | Intranuclear |
| HLADR | G46-6 | BV650 | T cell activation | Surface |
| CD39 | eBioA1 (A1) | FITC, PE Cy7 | T cell activation | Surface |
| CD25 | BC96, M-A251 | PE, BV605 | T cell activation | Surface |
| CD69 | TP1.55.3 | ECD | T cell activation | Surface |
| ki67 | B56 | AF647 | T cell proliferation | Intracellular |
| TIGIT | MBSA43 | FITC, PerCP EFluor710 | T cell activation/exhaustion | Surface |
| PD1 | EH12.2H7 | BV605 | T cell activation/exhaustion | Surface |
| IFNγ | 4S.B3 | FITC, BV510 | Cytokine | Intracellular |
| TNFα | Mab11 | AF700, PerCP Cy5.5 | Cytokine | Intracellular |
| CD107a | H4A3 | APC, BV605 | Degranulation | Surface |
| LIVE/DEAD | ---- | Near IR | Dead cell stain | ---- |

### Ex vivo analysis of MAIT function with partially fixed bacteria

MAIT cell function was measured *in vitro* by adapting assays that were previously described [25,26]. Briefly, cryopreserved PBMC were rested for 6 hours in RPMI (Thermo Fisher Scientific) supplemented with 10% FBS, 4 mM L-glutamine, and 1X antibiotic/antimycotic (Thermo Fisher Scientific Cat No. 15240062); this media will now be referred to as R10. For the stimulus, *Escherichia coli* and *M. smegmatis* stocks were fixed for approximately three minutes with a 1% paraformaldehyde solution, washed three times with 1X PBS, and diluted to a working concentration of 1×10^6^ CFU/mL in R10. After the 6-hour resting period, approximately 10 CFU of the indicated fixed bacterium were added for each PBMC cell. Cells were then incubated for 16 hours at 37°C in the presence of 2 uM monensin (Biolegend; San Diego, CA), 5 μg/mL Brefeldin A (Biolegend; San Diego, CA), and CD107a-APC (Table 1). Cells isolated from LN were incubated with bacteria for only 5 hours as the MAIT cell TCR is downregulated if the incubation period was too long (data not shown). After incubation, cells were stained with the Mamu MR1-5OPRU tetramer in R10+500 nM Dasatinib (Thermo Fisher Scientific) for 1 hour. Cells were washed once with 1X PBS (Thermo Fisher Scientific) and live cells stained using the LIVE/DEAD NearIR staining kit (Thermo Fisher Scientific) at a 1:4,000 dilution in 1X PBS. Cells were washed once with PBS containing 10% FBS and then stained with CD3, CD4, and CD8 antibodies (Table 1). Cells were fixed in 2% paraformaldehyde for 20 minutes, permeabilized in Medium B, and stained with IFN-γ and TNF (Table 1). Flow cytometry was performed as indicated above.

### Statistical analysis

Pearson’s correlation coefficients were calculated to determine the linear relationship between the absolute MAIT cell counts and the Log10 CFU within each granuloma. For comparative analysis of MAIT cell frequencies and phenotypes across cohorts, Mann-Whitney statistical tests were performed where indicated. For characterization of longitudinal changes across the same animals in a given cohort, non-parametric ANOVA tests were performed to determine statistical significance. In situations where not all animals had longitudinally matched data (Fig. 4B, C and 5D), Wilcoxon Rank signed tests were performed for matched pairs. To characterize longitudinal changes in the phenotypic markers present on MAIT cells during SIV infection, the data were first normalized to the average frequency for each marker before SIV infection for each animal. Then, the fold-change in expression of the indicated phenotypic marker relative to pre-infection measurement was calculated for every time point. Statistical significance was determined using ANOVA tests.

**Figure 1.**
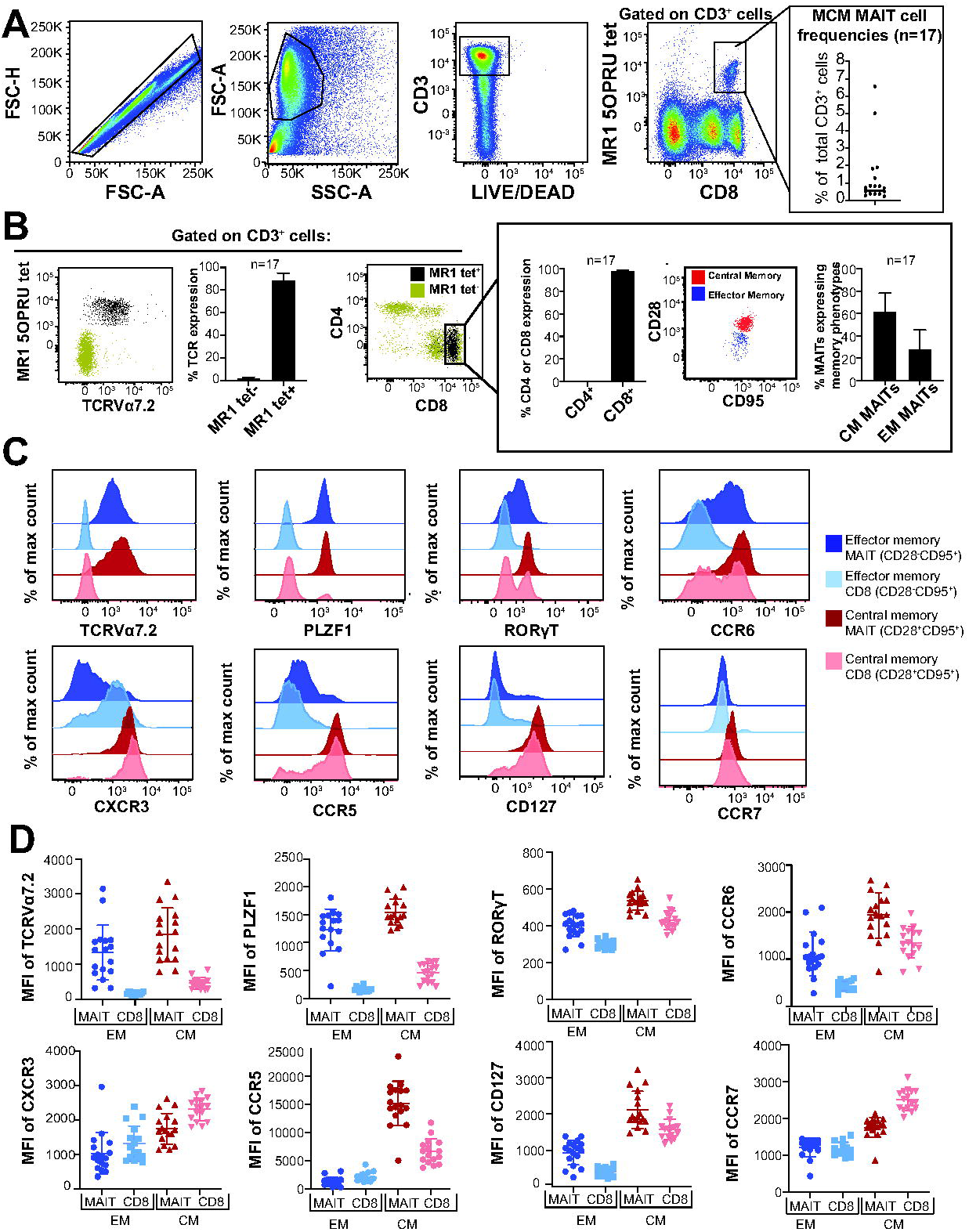
Characterization of MAIT cells in SIV-naïve MCM. A, Cryopreserved PBMC from 17 SIV naïve Mauritian Cynomolgus macaques (MCM) were stained with the MR1-5OPRU tetramer along with antibodies to CD3, CD4, CD8, TCRVα7.2, CD28, CD95, PLZF1, RORγT, CCR6, CXCR3, CCR5, CD127, and CCR7. Flow cytometry was performed as indicated in methods. Shown is a representative gating schematic to determine the frequency of CD3+MR1tet+ cells. The frequency of CD3+MR1tet+cells for all samples is shown. B, CD3+MR1tet+ (black dots) and CD3+MR1tet- (gold dots) cells were characterized for expression of CD4, CD8, and TCRVα7.2. Then, the frequency of MR1tet+TCRVα7.2+ cells from all animals that expressed CD4 or CD8 was determined. MR1tetramer+TCRVα7.2+ cells were characterized for central memory (CD28+CD95+, red dots) or effector memory (CD28-CD95+, blue dots) phenotypes, and the frequency of each memory subset for all MCM was determined. C and D, The indicated effector and central memory MAIT and bulk CD8 T cell populations were characterized for their expression of the indicated markers. Figure C shows a representative sample; Figure D shows the overall expression of the indicated markers across the population.

**Figure 2.**
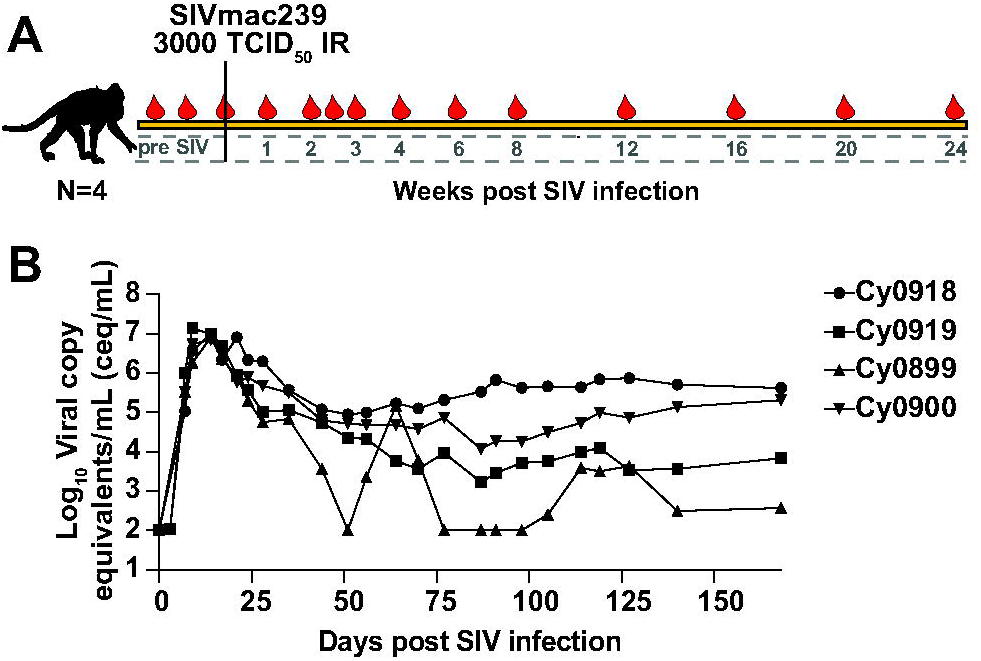
SIVmac239 infection of MCM. A, Four MCM were infected intrarectally with 3000 TCID50 of SIVmac239. Blood draw collection dates at the indicated times post-SIV infection are depicted by the red droplets. B, Plasma from the four SIVmac239-infected MCM indicated in (A) was collected and viral loads were determined as indicated in the methods. The Log_10_ virus copy equivalents/mL (ceq/mL) were graphed for each timepoint.

**Figure 3.**
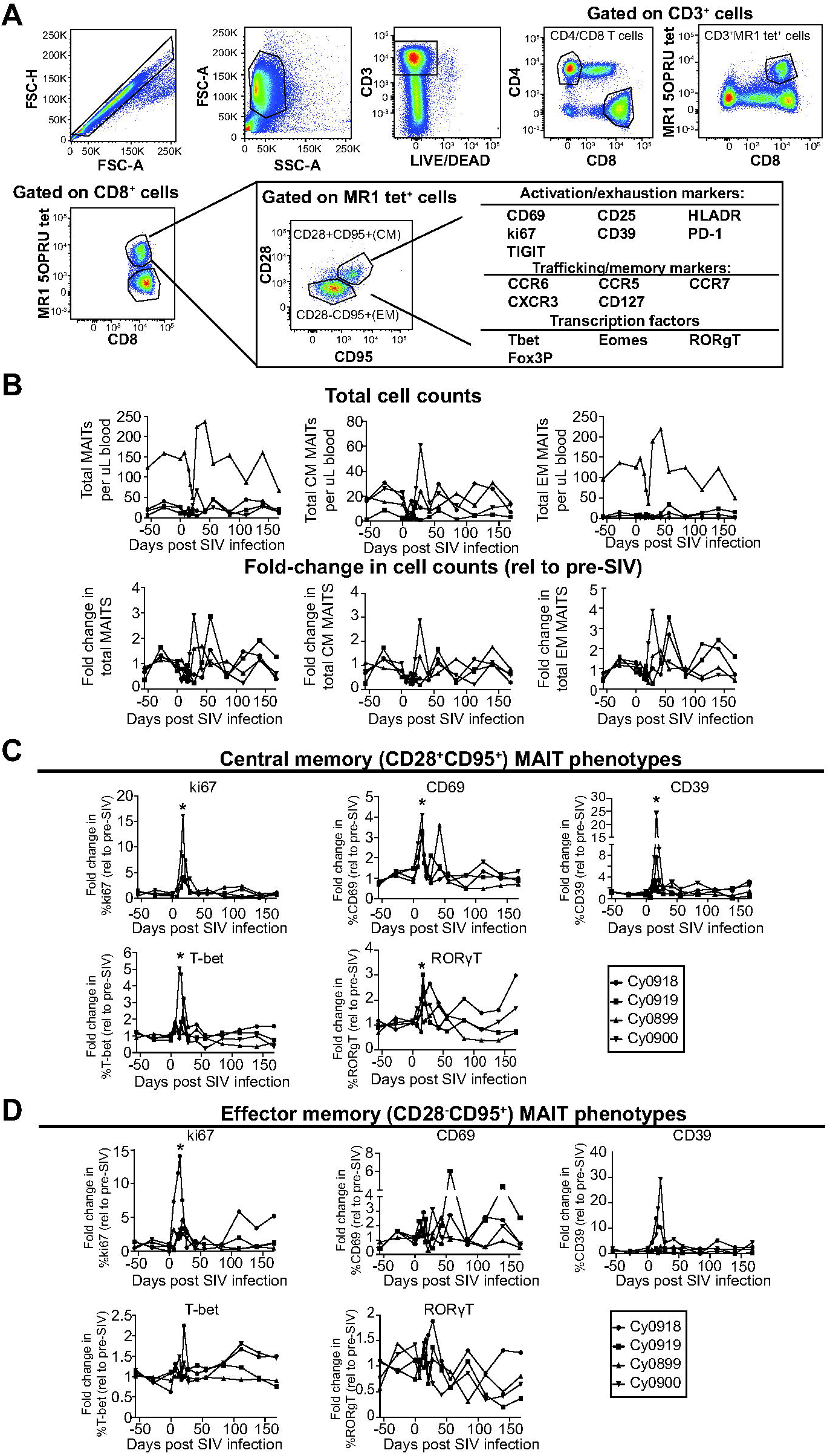
MAIT cells in the peripheral blood express an activated phenotype during acute SIV infection, but do not change in frequency longitudinally. A, Cryopreserved PBMC from the timepoints indicated in Fig. 2A were used and flow cytometry was performed. Shown is the gating schematic used to examine the expression of the indicated markers on the effector (CD28-CD95+, EM) and central (CD28+CD95+, CM) memory subpopulations within the CD8+MR1tet+ parent gate. B, Total CM and EM MAIT cell counts (top panels) were determined for each animal for each timepoint post-SIV infection based on complete blood counts measured on the day of blood collection. The fold-change in cell counts (bottom panels) was determined by dividing the total cell counts for each timepoint by the average of the total cell counts pre-SIV infection. C and D. Flow cytometry was performed as described in (A) and the fold change in the percent of CM (C) or EM (D) MAIT cells expressing the indicated markers relative to their pre-SIV expression levels were determined.

**Figure 4.**
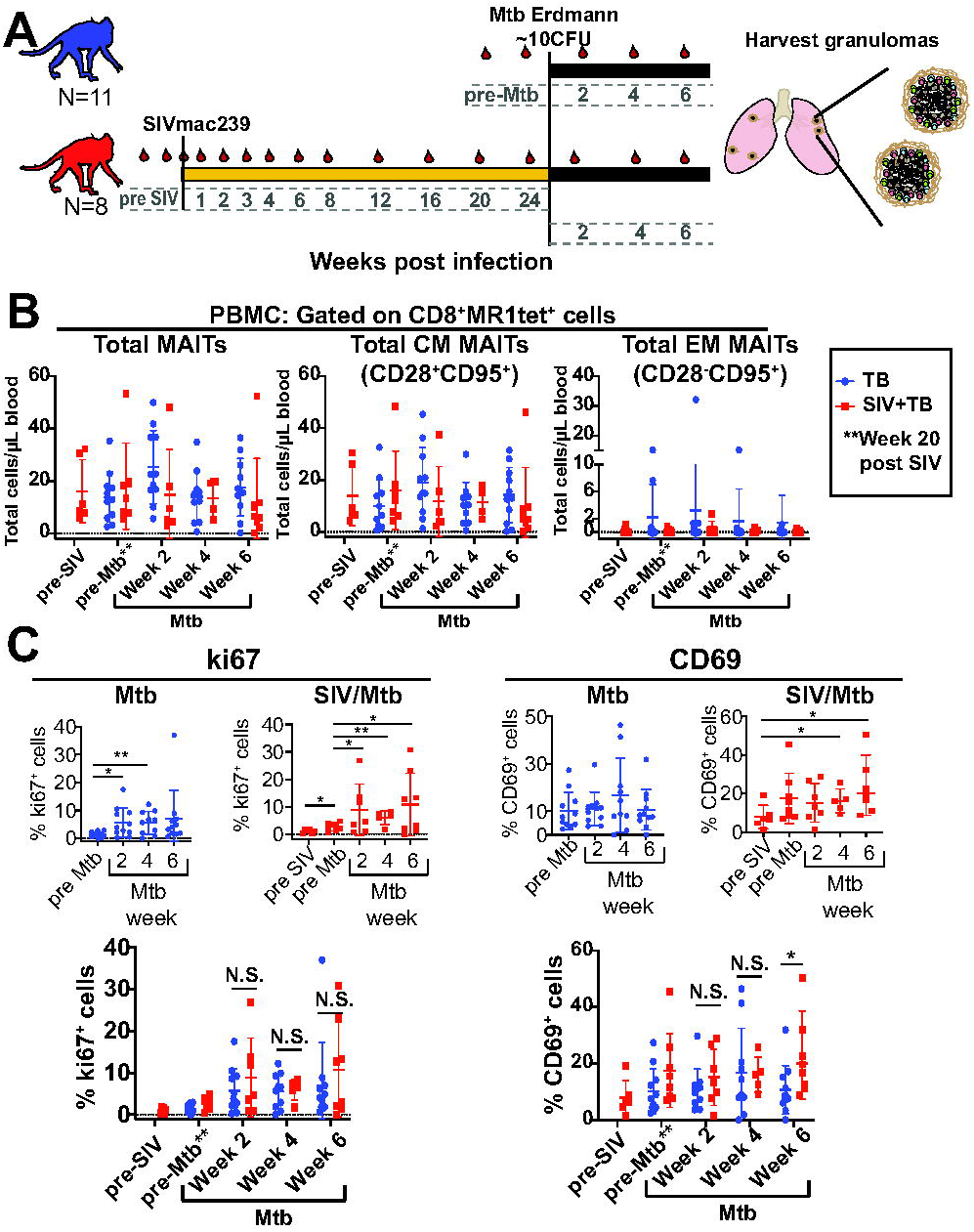
MAIT cells upregulate ki67 during Mtb infection in the peripheral blood, but do not increase in frequency during Mtb infection for SIV-naïve or SIV+ MCM. A, Schematic of the study design to investigate the role of MAIT cells during Mtb infection in SIV-naïve (blue, n=11) or SIV+ (red, n=8) MCM. Red droplets indicate blood collection timepoints. B, Cryopreserved PBMC from the indicated timepoints post-SIV and/or post-Mtb infection were thawed, stained for MAIT cells, and flow cytometry was performed as described in Fig. 3A. Total MAITs (left panel), central memory (CM) MAITs (middle panel), and effector memory (EM) MAITs (right panel) for SIV-naïve MCM (blue dots) or SIV+ MCM (red dots) were calculated based on complete blood counts collected at the time of the blood draw. C, Flow cytometry was performed as described in Fig. 3A to measure the expression of the indicated phenotypic markers on MAIT cells. The expression of ki67 and CD69 on MAIT cells was measured longitudinally for each cohort (top panels). Wilcoxon Rank signed tests for matched pairs were performed to determine statistical significance comparing pre-infection timepoints to post-infection timepoints; * indicates p<0.05, ** indicates p<0.005. Mann-whitney tests were used to compare differences between SIV-naïve (blue dots) and SIV+ (red dots) MCM for the same timepoints post-Mtb infection; * indicates p<0.05. N.S.=Not significant.

**Figure 5.**
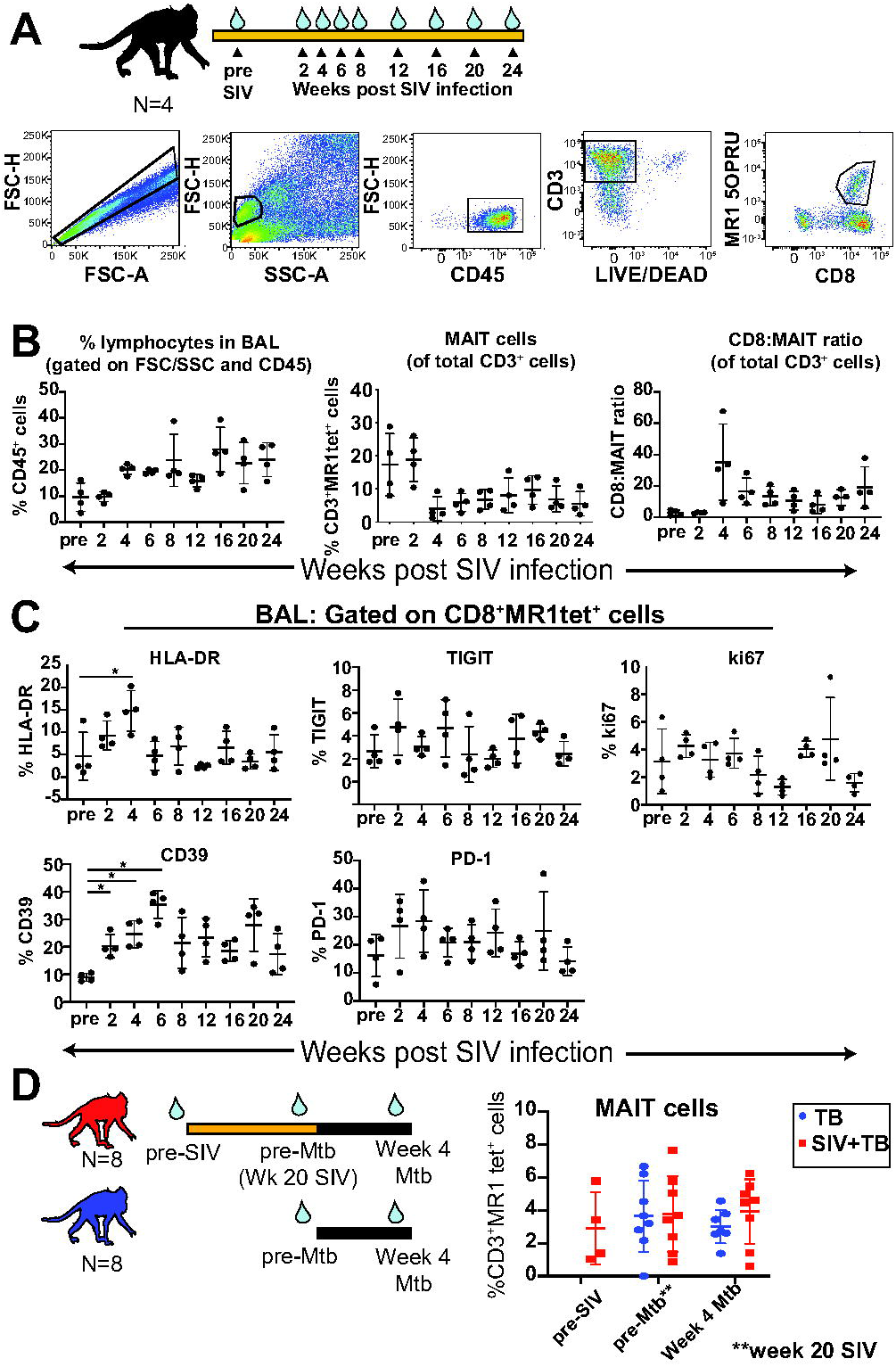
Longitudinal analysis of MAIT cells in bronchoalveolar lavage (BAL). A, BAL fluid from SIVmac239-infected MCM was collected at the indicated time post-SIV infection (arrows). Flow cytometry was performed and a representative image of the gating schematic used to determine MAIT cell frequencies within BAL is shown. B, The percent of lymphocytes present within each BAL sample for each indicated timepoint was determined (left panel). Within the CD3+ parent gate, the frequency of MR1tet+ cells (middle panel) and the CD8:MAIT cell ratio (right panel) were determined for each timepoint. C, The frequency of HLA-DR, TIGIT, ki67, CD39, and PD-1 was determined for theCD8+MR1tetr+ cells. ANOVA tests were performed to determine statistical significance between timepoints; *: p≤ 0.05. D, BAL was collected at the indicated timepoints (blue droplets) pre- and post-SIV and/or Mtb infection for 8 SIV/Mtb coinfected MCM (red) andr 8 Mtb-infected MCM (blue). Flow cytometry was performed as described in (A) and the frequency of CD3+MR1tet+ cells was determined for the indicated timepoints.

## RESULTS

### CHARACTERIZATION OF MAIT CELLS IN SIV-NAÏVE MCM

We first defined the phenotype of MAIT cells in SIV-naïve MCMs. Frozen PBMC from 17 SIV-naïve MCM were stained with the Mamu-MR1-5OP-RU tetramer. The MR1 tetramer-positive cells in the blood, henceforth referred to as MAIT cells, were then further characterized. The gating schematic for MAIT cells is shown in Figure 1A. MAIT cells comprised an average of 1.36% ±1.75% of the CD3+ T cells across the MCM tested (Fig. 1A). Nearly 100% of the circulating MAIT cells were CD4-CD8+ in all animals examined (Fig. 1B). MAIT cells comprised from 0.67% to 15% of the bulk CD8+ T cell population (data not shown).

The majority (>90%) of MAIT cells expressed the Vα7.2 T cell receptor (TCRVα7.2) [27], whereas <1% of MR1 tetramer-negative cells expressed TCRVα7.2 (Fig. 1B). We found that ∼61% ±16.48% of circulating MAIT cells expressed a central memory (CM) phenotype (CD28+CD95+) and that ∼28% ±17% expressed an effector memory (EM) phenotype (CD28-CD95+; Fig. 1B).

We also examined the expression of surface markers commonly found on MAIT cells, such as CCR6 [28], PLZF1 [29], and RORγT [28,30,31]. An example of the typical expression of these markers on the CM and EM subpopulations of MAIT and CD8+ T cells is shown in Fig 1C. Fig 1D shows the mean fluorescence intensity (MFI) of each marker on MAIT cells and CD8+ T cells across all MCM. Similar to previous studies [5,32,33], MAIT cells but not conventional CD8+ T cells expressed higher levels of TCRVα7.2 and PLZF1 (Fig. 1C-D). Also consistent with previous studies (reviewed in [34]), CM MAIT cells expressed high levels of RORγT and CCR6, whereas the expression of these two markers was lower on CM CD8+ T cells, EM MAIT cells, and EM CD8+ T cells (Fig 1C-D).

We examined the expression of CXCR3 and CCR5, both of which are associated with the trafficking of T cells to the lungs [35]. CXCR3 and CCR5 expression were very high on both CM MAIT cells and T cells compared EM MAIT cells (Fig 1C-D). We also measured expression of the LN homing marker CCR7 and the IL-7 receptor CD127 on MAIT and CD8+ T cells. Similar to other studies [36], we found that both CCR7 and CD127 were more highly expressed on CM MAIT and CD8+ T cells, when compared to EM MAIT and CD8+ T cells (Fig 1C-D).

### MAIT CELL NUMBERS ARE NOT REDUCED DURING 6 MONTHS OF SIV INFECTION

Several cross-sectional studies reported fewer MAIT cells in HIV- or SIV-infected individuals, when compared to healthy controls [8,10,12,13,15]. In contrast, a recent longitudinal study showed no decline in MAIT cell frequencies in SIV- or SHIV-infected pigtailed macaques [17]. Here, we infected 4 MCM intrarectally with 3,000 TCID_50_ of SIVmac239 and followed them for 6 months to determine if MAIT cells were depleted in the blood. (Fig. 2A). The viral load set point in the animals ranged from 10^2^ to 10^6^ viral copy equivalents/mL (Fig. 2B).

The frequencies and phenotypes of peripheral MAIT cells were measured longitudinally in these 4 SIV-infected animals (Fig 3A). We found that the frequency of MAIT cells was highly variable across the animals (Fig. 3B). Due to the large fluctuations in MAIT cell frequencies prior to SIV infection and the small number of animals, there were no statistically significant trends in total, CM, or EM MAIT cell frequencies in blood during the 6 months of SIV infection (Fig 3B), similar to what was observed in [17].

We then characterized the phenotypes of CM and EM MAIT cells during SIV infection using the markers indicated in Table 1 and gating strategy shown in Fig. 3A. We found a transient, but significant, increase in the expression of ki67, CD69, CD39, T-bet, and RORγT on CM MAIT cells between days 14-21 (Fig 3C, top panels). In contrast, EM MAIT cells had a statistically significant increase in only ki67 (Fig 3C, bottom panels) between days 14-21. While the activation markers CD69 and CD39 also showed transient increases, these changes did not reach the level of significance (Fig 3C). Overall, these data suggest that MAIT cells are activated during acute SIV infection (Days 14-21 post infection), but then returned to basal activation levels.

### SIV INFECTION DOES NOT IMPAIR CIRCULATING MAIT CELL FREQUENCY OR PHENOTYPE DURING MTB INFECTION

The role of MAIT cells within granulomas of Mtb-infected individuals is not well established and whether SIV co-infection alters MAIT cells within Mtb-affected tissues is unknown. Here, we used our SIV/Mtb co-infection model [18]to determine whether SIV disrupts the response of MAIT cells to Mtb infection. We bronchoscopically infected 11 MCM with a low dose of Mtb. Eight other MCM were first infected intrarectally with 3,000 TCID_50_ SIV for 6 months and then co-infected with the same dose and route of Mtb (Fig. 4A). Six weeks after Mtb infection, all animals were necropsied to collect granulomas and affected LN for pathology and flow cytometric analysis. At this relatively early time point post Mtb infection (or co-infection), the TB disease manifestation (e.g. pathology, bacterial load) did not differ significantly between the two cohorts, as further detailed in (Larson et al., manuscript in prep).

We examined the frequency of circulating MAIT cells within the two cohorts (Fig. 4B, left panel). Similar to the animals infected with SIV alone (Fig. 3), chronic SIV infection did not lead to differences in the absolute numbers of circulating MAIT cells (Fig. 4A, left panel; red dots). The absolute numbers of MAIT cells also remained unchanged for both the SIV-naïve and SIV+ group following infection with Mtb (Fig. 4B, left panel; blue dots vs. red dots). Additionally, no changes were observed in the absolute numbers of CM or EM subpopulations of MAIT cells (Fig. 4B).

Phenotypic changes in the expression of surface markers and transcription factors associated with T cell activation, proliferation, differentiation, and exhaustion were determined as described in Fig. 3A. While most phenotypic markers remained unchanged for both cohorts (data not shown), there were statistically significant increases in the frequency of circulating MAIT cells expressing ki67 following Mtb infection in both SIV-naïve and SIV+ animals (Fig 4C, left top panels). The increased frequency of MAIT cells expressing ki67 was statistically similar in both cohorts at all time points post-Mtb infection (Fig 4C, left bottom panel). Interestingly, the frequency of circulating MAIT cells expressing the early activation marker CD69 trended significantly higher following Mtb co-infection in SIV+ animals, but not in the SIV-naïve cohort (Fig 4C, right top panels). The frequency of CD69+ MAIT cells at 6 weeks post-Mtb infection was significantly higher in SIV+ MCM, compared to animals who were SIV-naïve (Fig 4C, right bottom panel).

### CHARACTERIZATION OF MAIT CELLS IN BAL DURING SIV AND/OR MTB INFECTION

BAL can be performed longitudinally following Mtb infection in macaques to assess the early immune response within the airways [37]. Additionally, the frequency and phenotype of MAIT cells within the BAL during SIV/SHIV [17] and Mtb [11] infection have been assessed. However, changes in MAIT cell frequency or function in the BAL following SIV infection and Mtb co-infection have not been assessed in MCM previously.

We collected longitudinal BAL samples during SIV infection and characterized the frequency and phenotype of MAIT cells as indicated in Fig. 5. At 4 weeks after SIV infection, there was an increase in the frequency of CD45+ lymphocytes within the airways, which remained elevated for the rest of the study (Fig. 5B, left panel). Concurrently, the percent of total airway CD3+ cells that were MR1 tetramer+ decreased (Fig. 5B, middle panel). This was likely related to an influx of CD3+CD8+MR1 tetramer-cells (Fig. 5B right panel, and Larson et al, manuscript in prep;), as previously reported for SIV infection[38–40], thus increasing the ratio of conventional CD8+ T cells to MAIT cells within the CD3+ parent gate.

Similar to the activation of circulating MAIT cells during SIV infection (Fig. 3), we also observed significant increases in MAIT cells expressing HLA-DR and CD39 in the BAL, peaking between 2-6 weeks after SIV infection (Fig. 5C, left-most panels). By 6 weeks after SIV infection, the frequency of MAIT cells expressing HLA-DR returned to pre-infection levels but the frequency of MAIT cells expressing CD39 remained elevated for the remainder of the study (Fig. 5C, left bottom panel). There were no changes during SIV infection in the frequency of MAIT cells expressing activation/exhaustion markers such as TIGIT and PD-1, or the proliferation marker, ki67 (Fig. 5C, middle and right panels).

We also collected BAL fluid from SIV+ and SIV-naïve animals after Mtb infection and performed flow cytometry (Fig. 5D). We found that Mtb infection did not affect the frequency of MAIT cells in either the SIV+ or SIV-naïve cohorts (Fig. 5D). The expression of activation markers on MAIT cells also was unaffected by Mtb infection in both cohorts (data not shown). Overall, these data show that airway MAIT cells are not affected by Mtb infection, irrespective of pre-existing SIV status.

### CHARACTERIZATION OF MAIT CELLS IN TISSUES

To assess whether a pre-existing SIV infection impairs MAIT cell frequency or function within Mtb-affected tissues, LN and granulomas were collected from SIV-naïve or SIV-positive MCM 6 weeks after Mtb infection (Fig. 4A) and flow cytometry was performed as indicated in Fig. 6A. The frequency of MAIT cells within Mtb-affected tissues was quantified for each cohort (Fig. 6B-C). Phenotypic differences in the expression of PD-1, TIGIT, ki67, IFN-γ, and TNF were measured on MAIT cells residing within the Mtb-affected tissues and compared between the two cohorts.

**Figure 6.**
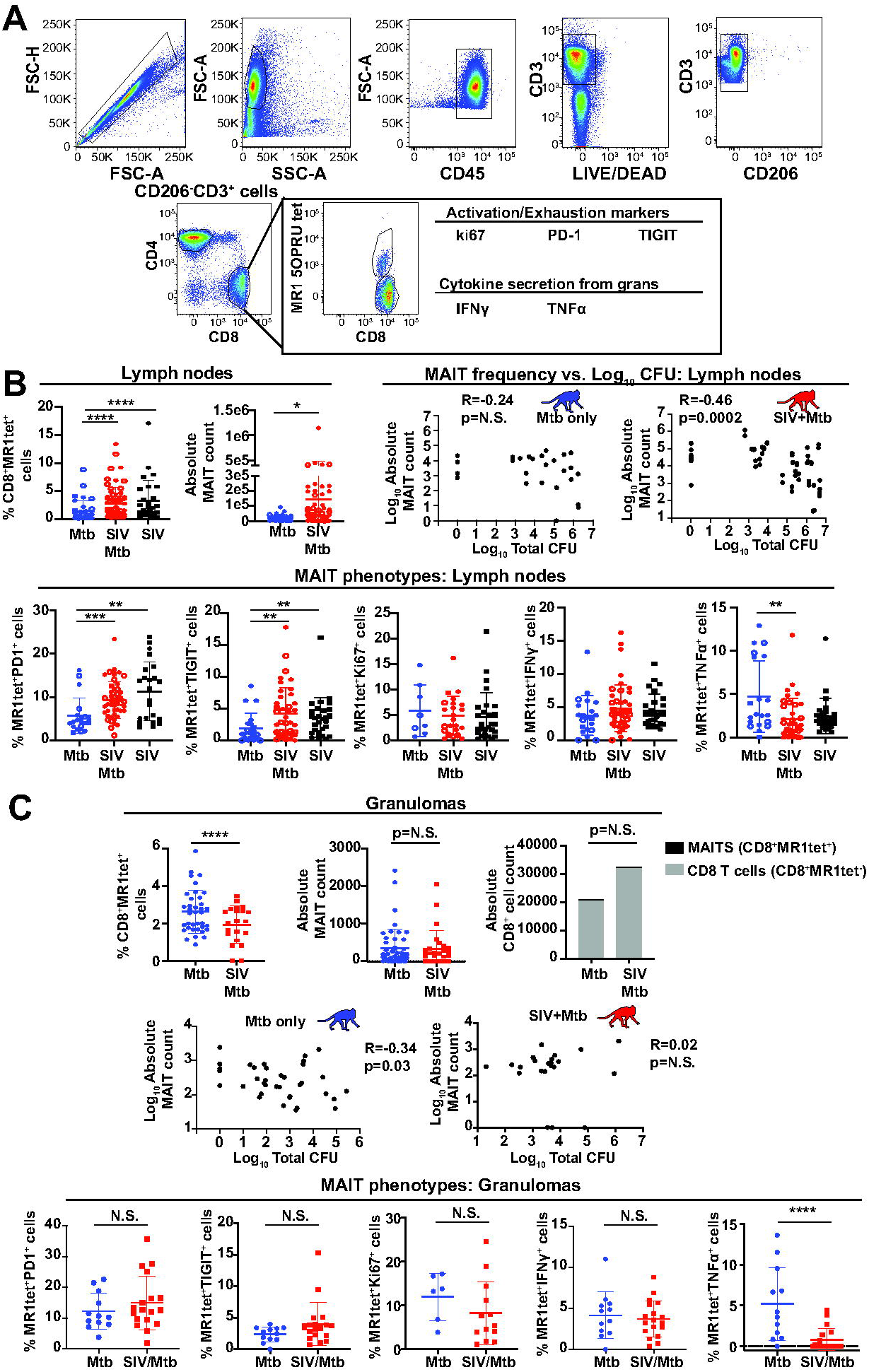
Analysis of MAIT cell frequency in Mtb-infected tissues. A, Mtb-infected tissues were collected 6weeks after Mtb infection from SIV-naïve and SIV+ MCM. Tissues were processed according to methods and cell homogenates were used for flow cytometric analysis. Shown is the gating schematic used to examine the expression of the indicated markers on the CD8+MR1tet+ cells. B, Homogenates collected from thoracic (circles) and extrathoracic (squares) lymph nodes present in MCM infected with Mtb (blue), SIV/Mtb (red), or SIV alone (black) were analyzed as indicated in (A) for the frequency of CD8+MR1tet+ cells. Open circles/squares indicate samples from animals infected with Mtb that were sterile. Absolute MAIT cell (CD8+MR1tet+) cell counts were determined based on total cell counts for individual samples. Mann-Whitney tests were performed to calculate statistical significance; ****: p<0.00005; *: p<0.05. Top right two panels: Pearson correlation coefficients were calculated to determine the relationship between the log-transformed values of the absolute MAIT cell counts and the log-transformed total colony forming units (CFU) for each sample. Shown are the correlations for samples collected from SIV-naïve (blue) and SIV+ (red). Bottom panels: the expression of the indicated markers were determined for CD8+MR1tet+ cells. Mann-Whitney tests were performed as indicated above for statistical significance; **: p<0.005, ***: p<0.0005. C, Top panels: CD8+MR1tet+ frequencies (left), and absolute cell counts (middle) for granulomas of Mtb-infected MCM (blue) and SIV/Mtb-infected MCM (red) were determined as in B. Top right; absolute CD8+ T cell (CD8+MR1tetr-; grey bars) and MAIT cell (CD8+MR1tet+, black bars) counts within granulomas were determined as described in (B). Middle panels: Pearson correlation coefficients were determined for the relationship between total MAIT cell counts and total CFU as indicated in (C). Bottom panels: MAIT cells (CD8+tetr+ cells) within granulomas were characterized for their expression of the indicated markers as described in (B). Mann-Whitney statistical tests were performed; ****:p<0.00005.

We found that within Mtb-affected LN, there was an increase in both the frequency and absolute numbers of MAIT cells in SIV/Mtb co-infected MCM compared to MCM infected with Mtb alone (Fig. 6B). This increase in MAIT cell frequency appeared to be related to SIV infection, as there was also a significantly increased MAIT cell frequency in peripheral LN of SIV+, but Mtb-uninfected, MCM (Fig. 6B). For both cohorts of Mtb-infected animals, we found weak negative correlations between MAIT cell frequency and bacterial burden (Fig. 6B, two right panels), although this correlation was not significant for the SIV-naïve group. MAIT cells in the LN of SIV+ animals, regardless of whether the animals were co-infected with Mtb, expressed higher levels of the activation/exhaustion markers PD-1 and TIGIT compared to SIV-naïve animals (Fig. 6B, bottom left panels). While the frequency of MAIT cells producing IFN-γ was not different between cohorts, we did observe a reduction in the frequency of MAIT cells producing TNF in SIV/Mtb co-infected MCM compared to MCM infected with Mtb alone (Fig. 6B, bottom right panel). Overall, we conclude that SIV infection leads to an increase in the frequency of activated MAIT cells in LN.

Similar comparisons were made for the cells isolated from granulomas of SIV+ and SIV-naïve animals following Mtb infection. The percentage of MAIT cells within the granulomas was lower in the SIV+ cohort, compared to the SIV-naïve cohort (Fig. 6C, top left panel). However, absolute MAIT cell counts remained similar between both cohorts (Fig. 6C, top middle panel). This suggests that an influx of non-MAIT CD8+ T cells into granulomas (Fig 6C, top right panel, and Larson et al., manuscript in prep) led to a decrease in the percent of MAIT cells in those sites. While there was a weak negative correlation between absolute MAIT cell counts and bacterial burden in the SIV-naïve cohort (Fig. 6C, middle left graph), there was no such correlation in the SIV+ cohort (Fig. 6C, middle right graph). In contrast to the LN, there were no differences in the frequencies of MAIT cells expressing PD-1, TIGIT, ki67, or IFN-γ (Fig. 6C, bottom 4 left panels) in the granulomas of SIV+ animals compared to those that were SIV-naïve. Notably, there was a significant decrease in the frequency of MAIT cells producing TNF in the SIV+ cohort compared to the SIV-naïve cohort (Fig. 6C, bottom right panel). Overall, the frequency of MAIT cells did not appear to correlate with bacterial burden from both SIV- and SIV+ animals. However, at the sites of Mtb infection, there was a decrease in TNF production in SIV+ MCM compared to SIV-MCM.

### EX VIVO ASSESSMENT OF MAIT CELL FUNCTION THROUGHOUT THE COURSE OF SIV INFECTION

MAIT cell function can be assessed ex vivo by using fixed bacteria as a stimulus [25]. However, despite the fact that mycobacteria-derived metabolites have been reported to activate MAIT cells[3,41], the vast majority of functional studies use *E. coli* as the bacterial stimulus [5,13,25,42]. Here, we tested the hypothesis that a mycobacterial stimulus would better reflect in vivo stimulation and reveal differences in MAIT cell function between the groups. PBMC from SIV-naïve MCM were incubated with 10 CFU/cell of fixed *E. coli* or *M. smegmatis* and flow cytometry was used to measure cytokine production and degranulation of MAIT cells in response (Fig. 7A). Incubation of PBMC with *E. coli* induced robust production of TNF and IFN-γ, as well as increased surface expression of the degranulation marker CD107a on MAIT cells (Fig. 7A, left 3 panels,). In contrast, there were no significant increases in the frequencies of MAIT cells producing CD107a, TNF, and IFN-γ from cells stimulated with *M. smegmatis* (Fig. 7A, left 3 panels) compared to unstimulated controls (Fig. 7A, left 3 panels).

**Figure 7.**
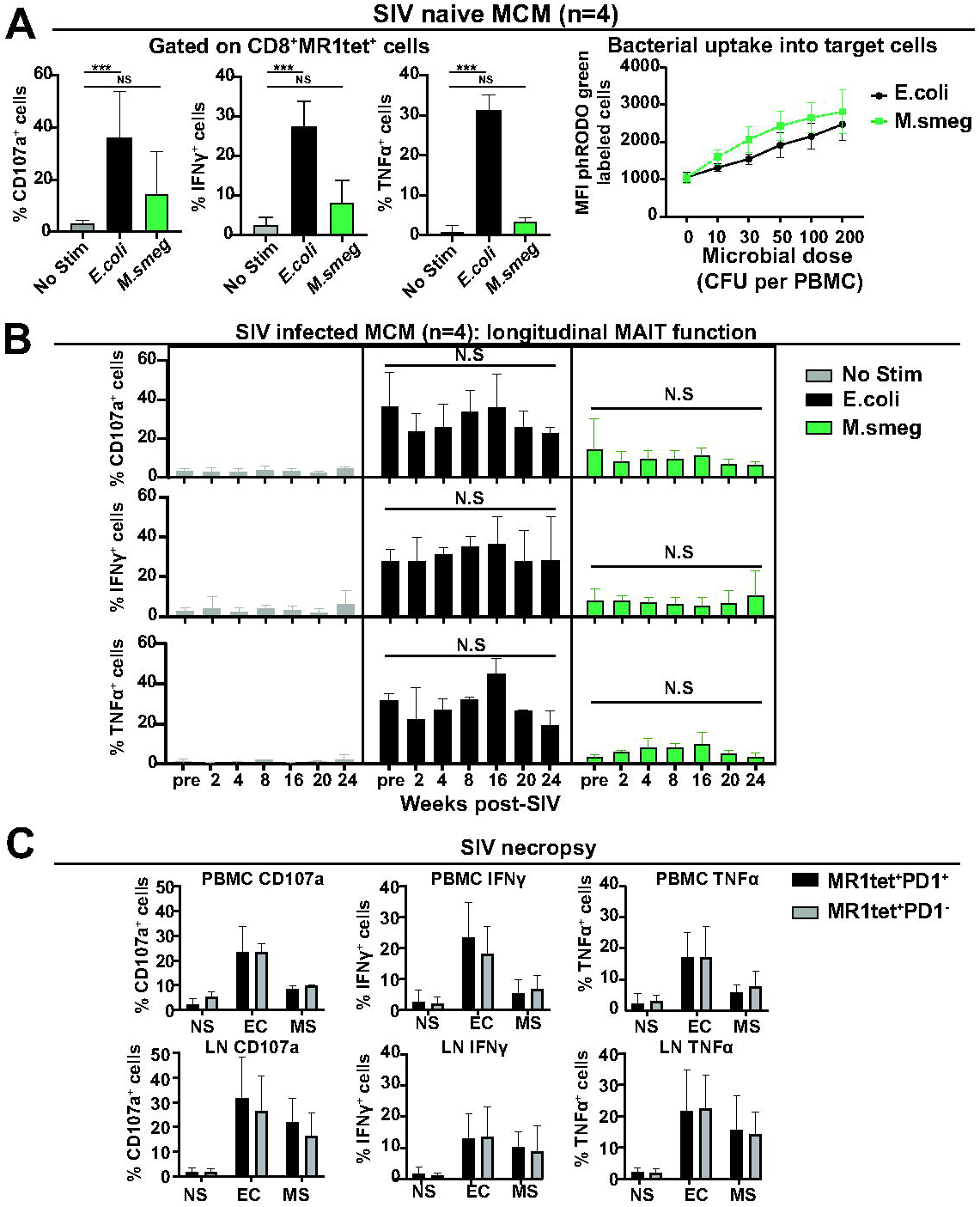
Longitudinal SIV infection does not impair ex vivo MAIT cell function. A, Cryopreserved PBMC from four SIV-naïve MCM were thawed, and functional assays were performed as described in the methods. Cells were stimulated for 16 hours with either 10 CFU of fixed *E.coli* (black bars) or *M.smegmatis* (green bars). The samples were stained as indicated in methods, and the expression of CD107a, IFN-γ, and TNF were measured on CD8+MR1tet+ cells from stimulated samples compared to an unstimulated sample control. Mann-whitney tests were performed to determine statistical significance; ***: p<0.0005. Right panel: Bacterial uptake into PBMC was measured by labeling equal amounts of bacteria with phRODO green dye as indicated in the methods, then incubating increasing microbial doses of the labeled bacteria with PBMC for 5 hours. B, Functional assays and flow cytometry were performed as described in (A) using cryopreserved PBMC collected at the indicated timepoints post-SIV infection for four SIVmac239 infected MCM. To compare expression of the indicated cytokines longitudinally for each individual animal for a single stimulus, ANOVA statistical analysis was performed. C, Cryopreserved lymph node (LN) and PBMC samples collected at the time of necropsy for the four SIVmac239 infected MCM described in (B) were used, and functional assays using fixed *E.coli* (EC) and *M.smegmatis* (MS), or no stimulation (NS) were performed as described in (A) and (B). MAIT cells were gated during flow cytometric analysis into two separate populations: CD8+MR1tet+PD1+ (black bars) and CD8+MR1tet+PD1- (gray bars). CD107a, IFN-γ, or TNF were measured as shown. Statistical analysis was performed as described in (A).

Previous work indicated that MAIT cells could react *in vitro* to *M. smegmatis*-derived metabolites [3,41]. Therefore, we wanted to determine whether the weak stimulation of MAIT cells with *M. smegmatis* we observed was related to poor uptake of the bacteria by antigen presenting cells. We labeled equal amounts of *M. smegmatis* and *E. coli* with a pHRODO green dye which increases in fluorescence intensity when labeled bacteria are taken up by antigen presenting cells. Increasing numbers of labeled bacteria were delivered to SIV-naïve PBMC and the fluorescence intensity was measured after 3 hours. For each dose of labeled bacteria tested, there were no significant differences in the ability of PBMC to uptake *M. smegmatis* compared to *E. coli* (Fig. 7A, right panel).

To determine if SIV infection alters MAIT cell function over time, frozen PBMC were incubated as indicated above with 10 CFU/cell of fixed *E. coli* or *M. smegmatis* (Fig. 7B). Despite the lack of reactivity of MAIT cells from SIV-naïve MCM to *M. smegmatis* (Fig. 7A), we continued to include this stimulus to determine if SIV and/or Mtb would increase MAIT cell responsiveness to *M. smegmatis*. We observed robust cytokine and CD107a production when cells were stimulated with *E. coli*, but weaker responses when *M. smegmatis* was used (Fig. 7B). No significant differences were observed in the production of TNF, IFN-γ, or CD107a at any time point measured pre- or post-SIV infection for either *E. coli* or *M. smegmatis* stimuli. Overall, these data suggest that SIV infection does not affect circulating MAIT cell function over time.

To determine if MAIT cell function was affected by the increased expression of PD1 or TIGIT we observed, functional assays were performed using PBMC and LN samples collected at the time of necropsy from animals infected with SIV alone. We found that in both PBMC and LN, cytokine production was similar for PD1+ and PD1-MAIT cells after stimulation with either *E. coli* or *M. smegmatis* (Fig. 7C). Taken together, no defects in ex vivo MAIT cell activity were apparent during SIV infection.

### REDUCED PRODUCTION OF TNF BY MAIT CELLS AFTER MTB INFECTION IN SIV+ MCM, COMPARED TO SIV-naïve MCM

While SIV infection alone did not affect MAIT cell function in response to ex vivo mycobacterial stimulus, we did observe an increase in the frequency of MAIT cells expressing ki67 in the blood for both Mtb and SIV/Mtb cohorts (Fig 4), indicating that MAIT cells may be proliferating and responding to an Mtb infection, as observed by others[7,11]. We performed *in vitro* functional assays using cryopreserved PBMC from the SIV-naïve and SIV+ animals following Mtb infection (Fig. 8). We observed robust cytokine production when PBMC from the Mtb-only animals were stimulated with *E. coli*, but not with *M. smegmatis* (Fig. 8A). There were no differences in MAIT cell cytokine production before or after Mtb infection. Likewise, for the SIV-coinfected MCM, MAIT cell function did not differ before or after SIV infection (Fig. 8B). However, there was a lower frequency of MAIT cells producing TNF after Mtb co-infection in response to both the *E. coli* and *M. smegmatis* stimuli (Fig. 8B, right panel). This is consistent with the reduced production of TNF by MAIT cells we observed directly ex vivo from LN and granulomas from SIV/Mtb co-infected MCM (Fig. 6).

**Figure 8.**
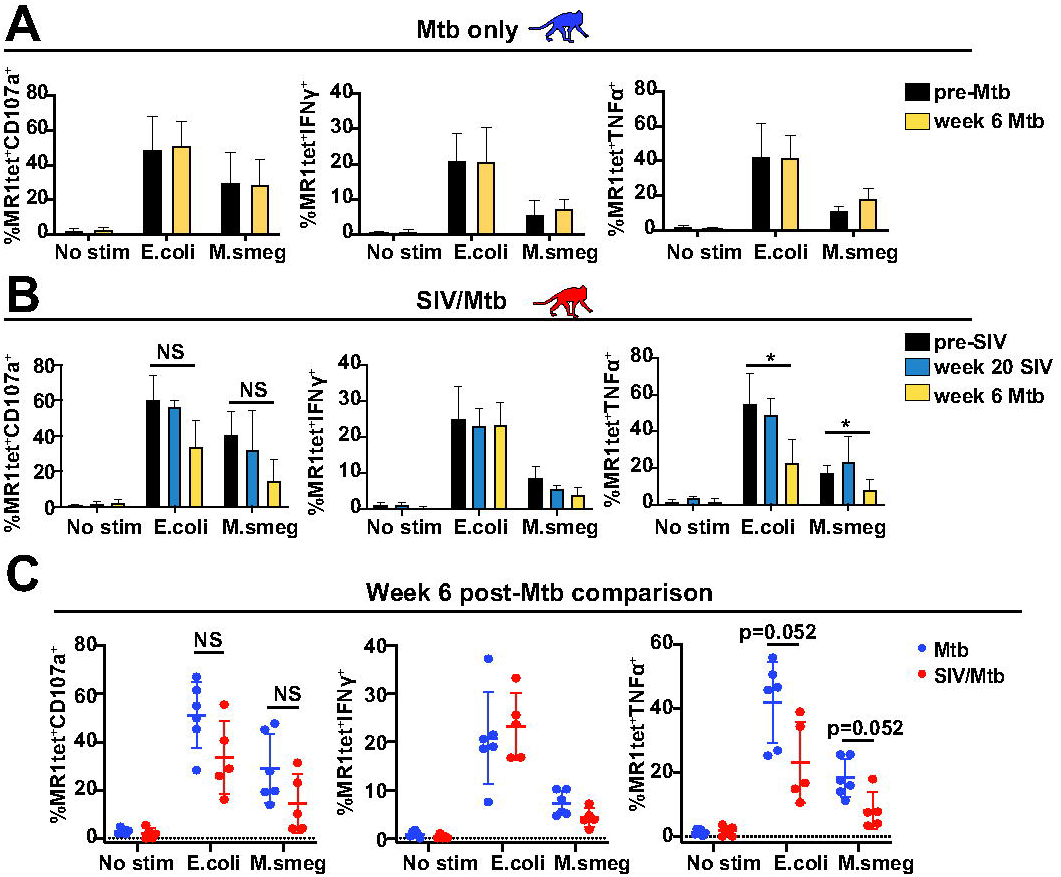
SIV and Mtb co-infection results in functional impairment of TNF production in MAIT cells in the peripheral blood. A, Cryopreserved PBMC from pre-Mtb (dark grey) and 6 weeks post-Mtb (yellow) infection from seven Mtb-infected MCM (blue monkeys) were thawed and functional assays were performed as described in Fig. 7A. ANOVA statistical tests were performed to compare the expression of the indicated cytokines or CD107a longitudinally for a single stimulus. B, Cryopreserved PBMC from pre-SIV (dark grey), Week 20 SIV (teal), and 6 weeks post-Mtb (yellow) infection from six SIV/Mtb coinfected MCM (red monkeys) were thawed and functional assays were performed as described in (A). ANOVA statistical tests were performed to compare the expression of the indicated cytokines and CD107a longitudinally for a single stimulus. C, The functional assays performed in (A) and (B) using frozen PBMC from six weeks post-Mtb infection for SIV-naïve (blue dots) and SIV+ (red dots) MCM were plotted on the same graphs to compare the expression of CD107a (first panel), IFN-γ (second panel), or TNF(third panel) amongst the two cohorts. Mann-whitney tests were performed to determine statistical significance.

## DISCUSSION

Here, we characterize in detail MAIT cells in the SIV, Mtb, and SIV/Mtb co-infection MCM model. We phenotype MAIT cells longitudinally during SIV infection, and also describe the first study in MCM focusing on the early MAIT cell response to Mtb infection within Mtb-affected tissues (e.g. granulomas). Finally, we show for the first time that SIV co-infection affects MAIT cell frequency and function within granulomas.

Notably, we showed a decrease in TNF production by MAIT cells from SIV/Mtb co-infected MCM compared to those from MCM infected with Mtb alone and was observed in both LN and granulomas (Fig. 6). Furthermore, although the decrease in TNF production was not statistically significant at 6 weeks post-Mtb infection between the SIV-naïve and SIV+ cohorts during ex vivo assays (Fig. 8C), there was a significant decline in ex vivo TNF production over time in the SIV+ animals following Mtb co-infection (Fig. 8B). This is consistent with the decreased TNF production we also observed from conventional CD4+ and CD8+ T cells specifically during SIV/Mtb co-infection (Larson et al, manuscript in prep) and suggests that defective production of TNF from T cells, both conventional and nonconventional, may play a key role in the profound susceptibility to Mtb in SIV+ macaques. The mechanism by which this defect in TNF production occurs has not yet been determined. It has been established that anti-TNF therapy or blocking TNF production can lead to exacerbation of primary TB or reactivation of latent Mtb infection in several animal models as well as Mtb-infected individuals [43–45] [46] [47]. In the study reported here, we performed necropsies just 6 weeks following Mtb infection (or co-infection) in order to capture early events post Mtb. There were no significant differences in measures of TB disease (e.g. pathology, PET/CT imaging, bacterial load) between the SIV+ or SIV-naïve MCM at this early time point. However, we showed previously that SIV+ MCM exhibit significantly more TB disease than SIV-naïve MCM when animals are followed for 8 weeks or more after Mtb infection [18]. The decline in MAIT cell TNF production we report here might lead to TB disease exacerbation if this TNF defect continues to become more pronounced as the Mtb infection progresses. Defects in TNF production also have been observed in humans during HIV infection and impair the ability of immune cells to respond to Mtb infection [48]. Therefore, reduced production of TNF from T cells in our MCM co-infection model could explain the increased TB disease burden previously observed in SIV+ macaques [18].

Another remarkable finding was the absence of MAIT cell responses to mycobacterial infection. There was no Mtb-mediated MAIT cell activation or proliferation within the granulomas (Fig. 6C) even though conventional CD4+ and CD8+ T cells in granulomas upregulated both activation and proliferation markers (Larson et al. manuscript in prep). Our findings are similar to a study performed in rhesus macaques that also showed that MAIT cells from granulomas did not respond to Mtb [11]. Further confirming our findings, we found that circulating MAIT cells could not produce cytokines or express CD107a when stimulated ex vivo with *M. smegmatis* (Fig.7A), while MAIT cells stimulated with *E. coli* robustly responded (Fig. 7A). Neither SIV nor Mtb infection improved the ability of MAIT cells to react to *M. smegmatis* (Figs. 7 and 8). One possible explanation for the findings reported here (Fig. 7 and 8) and by others [11] is that the T cell receptors present on macaque MAIT cells are not well-suited for recognizing mycobacterial metabolites. MAIT cells do display ligand discrimination depending on their TCR usage [41], and so it is possible that the predominant TCRs used by MAIT cells within the lungs of macaques do not react as robustly to mycobacterial ligands. Other lines of evidence support that MAIT cells may not respond robustly to mycobacteria as PBMC incubated with MR1 tetramers made using metabolites from *E. coli* resulted in more cells producing IFN-γ than in PBMC incubated with tetramers incorporating *M. smegmatis* metabolites [49]. Overall, our findings support those of others [49] that MAIT cells within the lungs of macaques may not respond appropriately to control Mtb infection and limit TB progression.

While it is unclear whether MAIT cells have a direct role in antimycobacterial immunity, SIV co-infection clearly affects their ability to secrete certain cytokines such as TNF (Fig. 8). Therefore, this dysregulation of MAIT cells may indirectly impair the antimycobacterial response of other effector cells. MAIT cells secrete several proinflammatory cytokines in response to bacterial infection [34]. Recently, it was shown that MAIT cells play critical roles in providing signals to conventional T cells which, in the context of Mtb infection, could activate them or direct them to the lungs or other sites of infection [50,51]. Furthermore, MAIT cells have recently been shown to provide signals to B cells as well to develop antibody responses to pathogens [52]. Whether SIV affects the ability of MAIT cells to recruit or activate other immune cells during Mtb infection has not been assessed, and may be interesting to pursue in future studies.

Another interesting observation was the phenotypic differences between MAIT cells within the LN versus those within granulomas. Granuloma-resident MAIT cells did not display any phenotypic differences between the SIV-naïve and SIV+ cohorts other than lower TNF production in the latter group. In contrast, the MAIT cells within Mtb-affected LN of SIV+ MCM displayed increased expression of PD-1 and TIGIT compared to SIV-naïve MCM (Fig. 6). This appeared to be driven by SIV infection specifically, as the MAIT cells within the LN of both SIV-infected (black dots) and SIV/Mtb co-infected (red dots) MCM expressed higher levels of these markers (Fig. 6B). While we did not find any evidence that these markers were associated with MAIT cell exhaustion (Fig. 7C), it is clear that ongoing SIV replication can lead to the activation of MAIT cells. This was also observed in the blood, where circulating MAIT cells briefly increased their expression of ki67, CD69, CD39, RORγT, and T-bet, coincident with peak viremia (Fig. 2, Fig. 3B and C). The increase in ki67 is consistent with increased MAIT cell proliferation observed by Juno and colleagues during acute SIV infection [17]. TCR-independent activation of MAIT cells has been observed for other viral infections as well [53]. We hypothesize that during both acute SIV infection and active SIV replication within the LN, other immune cells are producing cytokines such as IL-12 and IL-18 [54–56]. These cytokines are known to activate MAIT cells [57] and, therefore, MAIT cell activation may be occurring indirectly though TCR-independent mechanisms. This hypothesis remains to be tested.

Overall, our findings demonstrate the absence of a direct role for MAIT cells in the antimycobacterial response within Mtb-infected MCM, similar to the findings in rhesus macaques [11]. Importantly, while MAIT cells do not respond robustly to mycobacterial stimulus ex vivo (Figs. 7-8), SIV co-infection does alter the phenotypic characteristics of MAIT cells at sites of Mtb infection (Fig. 6) and impairs their ability to produce pro-inflammatory cytokines such as TNF (Fig. 8). As discussed above, the role that MAIT cells play in antimycobacterial immunity may be indirect, and the phenotypic changes induced during SIV infection may be at least partially responsible for the deleterious effects of Mtb in HIV/SIV co-infected individuals.

